# Evaluating brain cell marker genes based on differential gene expression and co-expression

**DOI:** 10.1101/554626

**Authors:** Rujia Dai, Yu Chen, Chuan Jiao, Jiacheng Dai, Chao Chen, Chunyu Liu

## Abstract

Reliable identification of brain cell types is necessary for studying brain cell biology. Many brain cell marker genes have been proposed, but their reliability has not been fully validated. We evaluated 540 commonly-used marker genes of astrocyte, microglia, neuron, and oligodendrocyte with six transcriptome and proteome datasets from purified human and mouse brain cells (n=125). By setting new criteria of cell-specific fold change, we identified 22 gold standard marker genes (GSM) with stable cell-specific expression. Our results call into question the specificity of many proposed marker genes. We used two single-cell transcriptome datasets from human and mouse brains to explore the co-expression of marker genes (n=3337). The mouse co-expression modules were perfectly preserved in human transcriptome, but the reverse was not. Also, we proposed new criteria for identifying marker genes based on both differential expression and co-expression data. We identified 16 novel candidate marker genes (NCM) for mouse and 18 for human independently, which have the potential for use in cell sorting or other tagging techniques. We validated the specificity of GSM and NCM by in-silico deconvolution analysis. Our systematic evaluation provides a list of credible marker genes to facilitate correct cell identification, cell labeling, and cell function studies.

## Introduction

The human brain is a heterogeneous organ with numerous cell types. It has billions of cells including half neurons and half glia^1^. The major classes of glia are astrocyte, microglia and oligodendrocyte. Identifying these cell types is important because it would permit the brain to be understood in greater detail and would be especially useful for studying cellular contributions to the psychiatric disorders. A critical need in neuroscience research, is to develop methods to reliably identify specific brain cell types.

A strategy that has been employed to identify specific cell types is the development of marker genes, which are sets of genes that express specifically in a cell type. Thousands of genes have been proposed as marker genes^2^. One well-known marker gene, RBFOX3 (gene of NeuN), is only expressed in nuclei of most neuronal cell types^3^. Marker genes can be used in several applications. Protein products of marker genes can be used to label different cell types, which may be used in fluorescence activated cell sorting (FACS). Marker genes also can be used to determine cell composition in bulk tissue samples. A computational method known as supervised deconvolution was developed to infer cell proportions in bulk tissue samples based on the expression of marker genes^4-6^. This method has been applied to studying the composition of bulk brain samples^7,8^. High specificity of marker genes is critical for generating reliable results in all of these applications.

Differential gene expression (DGE) analysis of transcriptome or proteome data is the most straightforward way to define the specificity of marker genes^9-15^. One of the drawbacks of DGE is that the outcomes is study-dependent. The outcomes are affected by many factors such as species, cell or tissue source, and the data generation platform. Human and mouse genomes are 80% orthologous^16^, but differences in gene expression between species are often greater than those between tissues within one species^17^. Within a species, cells isolated from primary culture or acutely from tissue showed different gene expression patterns^18^. Also, the expression estimates of the marker may vary considerably depending on whether mRNA or protein is measured. The statistical variation in transcriptome only explained 40% of the statistical variation in protein level^19^. Besides these biological confounders, the experimental platforms used to quantify gene expression level may also impact marker gene selection. RNA-Seq provides a larger dynamic range for the detection of transcripts and has less background noise, resulting in RNA-Seq being more sensitive in calling cell type-specific genes than microarray platforms^20^. Another weakness of DGE is that relationships among marker genes are not considered in the analysis. Groups of marker genes are often used to describe a cell type, and marker genes work with each other to execute functions in specific cell type. The relationship between marker genes represents their coordinated functions, specificities, and expressions. In DGE analysis, marker genes are defined independently, and the relationship among them is ignored.

Co-expression (COE) is a method of identifying interactions among genes by assigning genes with similar expression patterns into a module^21,22^. There was study reported that the co-expression modules in brain enriched cell type marker genes^23^. So it suggested that the co-expression can detected the cell type-specific marker genes, even in the heterogenous samples. The module formed by marker genes indicates their coordinated functions and specificities for a cell type. The correlation of genes with cell type-specific module suggests it’s cell specificity. COE has the potential to systematically capture marker genes group that DGE cannot.

In this study, we evaluated the specificity of 540 published brain cell marker genes and discovered novel marker genes by DGE and COE analyses. We used six datasets containing transcriptome and proteome data from purified astrocytes, microglia, neurons and oligodendrocytes from both mouse and human brains. We identified 22 brain cell marker genes out of the 540 candidates, referred as gold-standard marker genes (GSM), that specifically express in one cell type. We constructed brain cell-related gene co-expression modules for human and mouse, and found large differences among species. We found a statistically significant correlation between cell-specific fold change, a measure developed in this study, and gene membership in the brain cell-related coexpression modules. Combining DGE and COE, we identified 16 novel candidate marker genes (NCM) in mouse brain and 18 NCM in the human brain. Through supervised cell deconvolution analysis, we showed that using GSM and NCM improved the performance of deconvolution.

## Results

To evaluate and discover brain cell marker genes, we performed DGE and COE analysis on transcriptomic or proteomic data (Figure 1). We used six datasets of purified cell populations for DGE analysis (DGEDat) and two single cell datasets for COE analysis (COEDat) (Table 1). The DGEDats included transcriptome and proteome data from human and mouse brain purified cell populations. The COEDats were single-cell RNA sequencing data from both human and mouse brains.

**Table 1.**
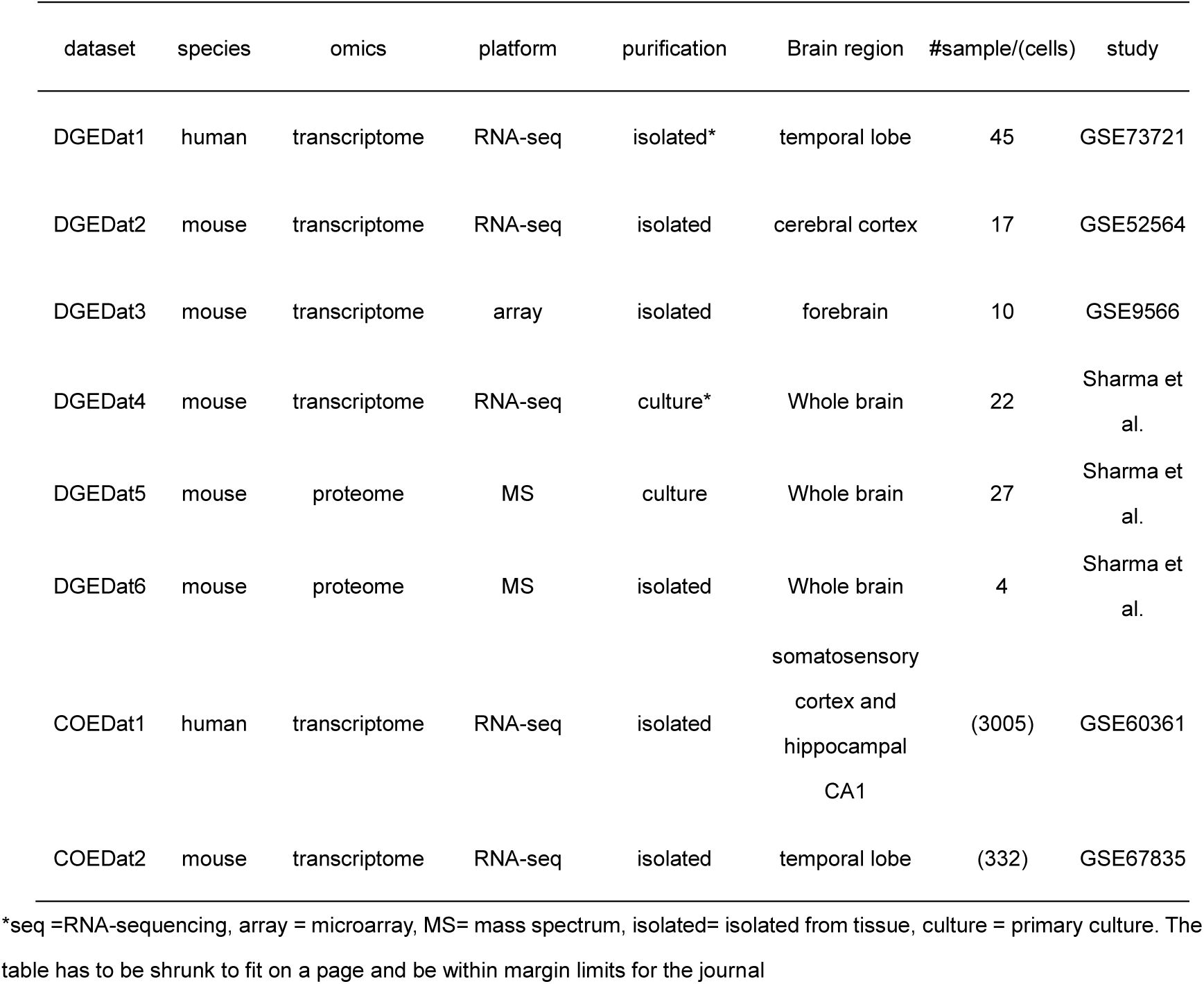
Datasets used

| dataset | species | omics | platform | purification | Brain region | #sample/(cells) | study |
| --- | --- | --- | --- | --- | --- | --- | --- |
| DGEDat1 | human | transcriptome | RNA-seq | isolated* | temporal lobe | 45 | GSE73721 |
| DGEDat2 | mouse | transcriptome | RNA-seq | isolated | cerebral cortex | 17 | GSE52564 |
| DGEDat3 | mouse | transcriptome | array | isolated | forebrain | 10 | GSE9566 |
| DGEDat4 | mouse | transcriptome | RNA-seq | culture* | Whole brain | 22 | Sharma et al. |
| DGEDat5 | mouse | proteome | MS | culture | Whole brain | 27 | Sharma et al. |
| DGEDat6 | mouse | proteome | MS | isolated | Whole brain | 4 | Sharma et al. |
| COEDat1 | human | transcriptome | RNA-seq | isolated | somatosensory cortex and hippocampal CA1 | (3005) | GSE60361 |
| COEDat2 | mouse | transcriptome | RNA-seq | isolated | temporal lobe | (332) | GSE67835 |

\*seq =RNA-sequencing, array = microarray, MS= mass spectrum, isolated= isolated from tissue, culture = primary culture. The table has to be shrunk to fit on a page and be within margin limits for the journal

**Figure 1.**
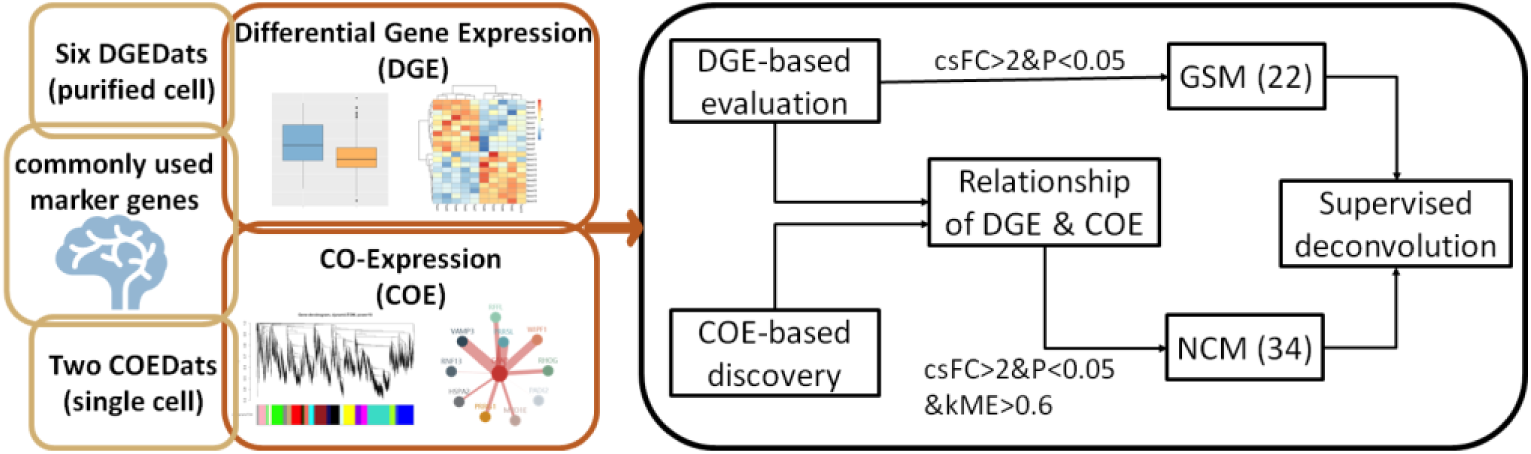
Analysis workflow. Six DGEDats of the purified cell population and two COEDats of single cells were used to evaluate 540 commonly-used brain cell marker genes. Differential gene expression (DGE) was performed on six DGEDats and the cell-specific fold change (csFC) was defined to measure the cell specificity for the marker genes. Co-expression (COE) analyses were performed on two COEDats and cell-specific networks were constructed. The correlation of genes with the module eigengene in the cell network was measured as module membership (kME). Through DGE-based evaluation, 22 gold-standard marker genes (GSM) were identified. Combining DGE and COE, 34 novel candidate marker genes (NCM) were identified. The specificities of GSM and NCM were demonstrated in supervised deconvolution.

### Commonly-used marker genes of four major cell types

We collected 540 marker genes that were commonly used for labeling cells and validating cell isolation (Supplementary Table 1). These marker genes were identified in published literature^9,10,13-15^, company websites^24,25^, and ISH databases, such as the Allen Brain Atlas (ABA) and GENSAT_^26-28^_ for labeling neurons, astrocytes, microglia, oligodendrocytes, and other cell types in the brain. Of 540 candidate marker genes, only eight genes were reported in all data sources while most of the marker genes were source-specific (Supplementary Figure 1). Genes annotated as marker genes of more than two cell types by different sources were considered as “conflict marker genes.” We found 27 conflict marker genes in the 540 collected genes (Supplementary Table 1). The other genes had no conflict annotations in different data sources and were classified as “consistent marker genes.”

### DGE-based specificity evaluation of commonly-used marker genes

We identified Gold-Standard Marker genes (GSM) that showed cell-type specificity across multiple types of data through DGE analysis. We found that the classical fold-change value, which is typically calculated as the expression in the target cell divided by averaged expression in other cells^14,29^, may produce inaccurate calls of marker genes (Supplementary Figure 2, Supplementary Table 2). To avoid this problem, we created a measure of cell-specific fold change (csFC). The csFC was defined as equation (1).

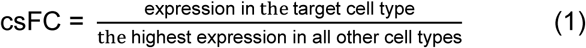

To be considered a GSM, the following four criteria had to be met based the datasets we collected: 1) the gene must be detected in the target cell type in all six DGEDats. There were 113 of the 540 candidates that met this criterion. 2) csFC ≥ 2 in all six DGEDats. 3) Benjamin-Hochberg (BH) corrected p-value of the two-sample Wilcoxon test of expression in the target cell, and expression in other cell types should be lower than 0.05 in more than two of the six DGEDats. 4) the gene must be shown to be specific in at least one proteomic dataset. Using these criteria, we identified 22 GSM in total. Nineteen of the 22 GSM were from the consistent marker genes group, and three were from the conflict marker genes group (Table 2).

**Table 2.**
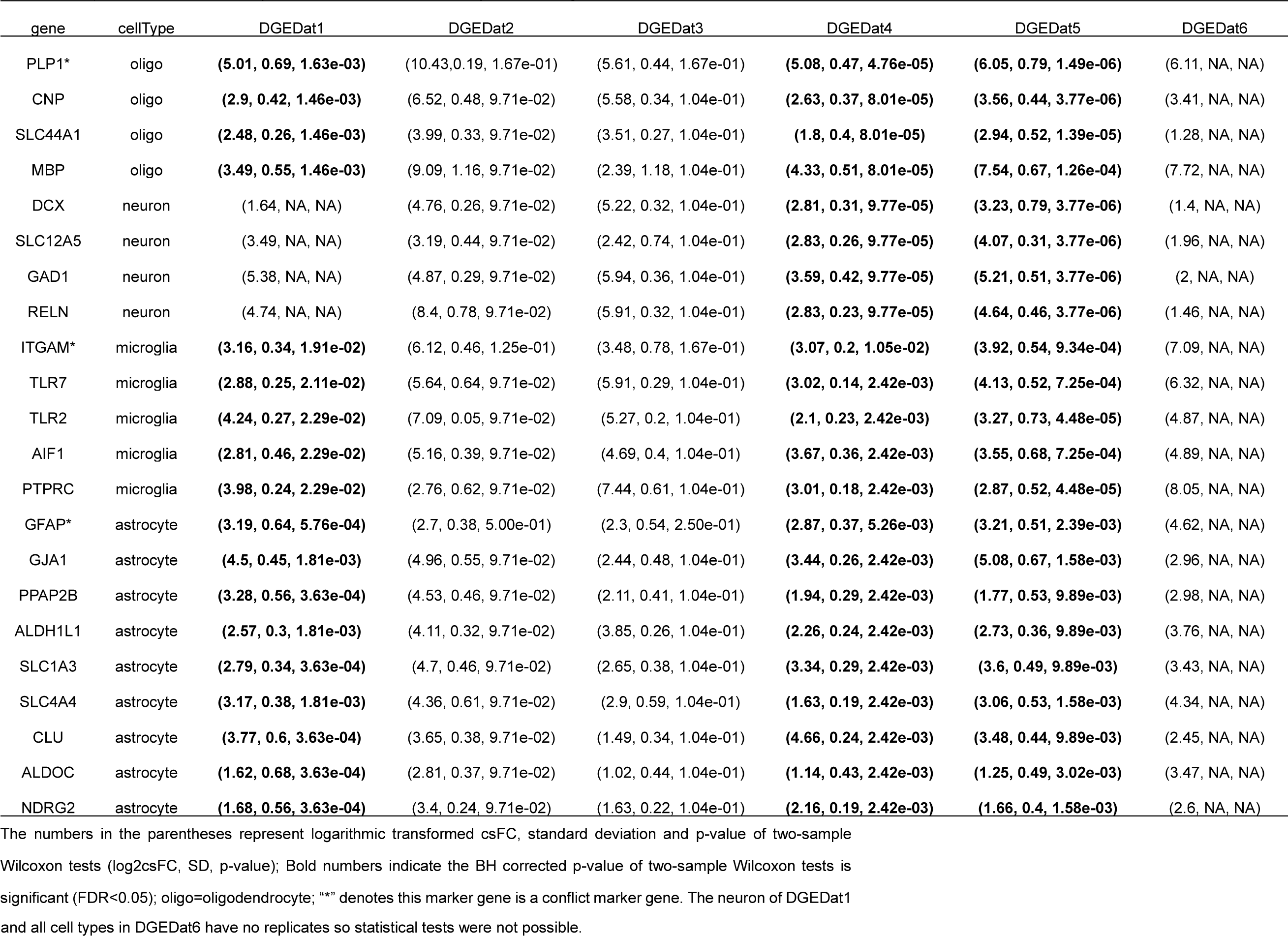
csFC, standard deviation and p-value of GSM in differential expression analysis of six DGEDats

### COE analysis of two large single-cell datasets

To discover the co-expression of marker genes, we performed weighted gene co-expression network analysis (WGCNA) on human and mouse brain single-cell transcriptome data in parallel with DGE. We annotated the co-expression modules using pSI packages^30^, which can identify genes enriched in specific cell populations and test gene overrepresentation by Fisher’s exact test. Figure 2A shows the p-value of cell type enrichment of each module after correcting for multiple testing by BH. We chose the most significant module in the cell type enrichment analysis as the brain cell co-expression module (BCCM) for each cell type (Table 3,Supplementary Figure 3 and Supplementary Figure 4). We used Gene Ontology analysis to determine the biological functions of each BCCM (Supplementary Table 3). The BCCMs were enriched in biological processes for specific cell types. For example, the oligodendrocyte-related module was enriched in the axon ensheathment pathway.

**Figure 2.**
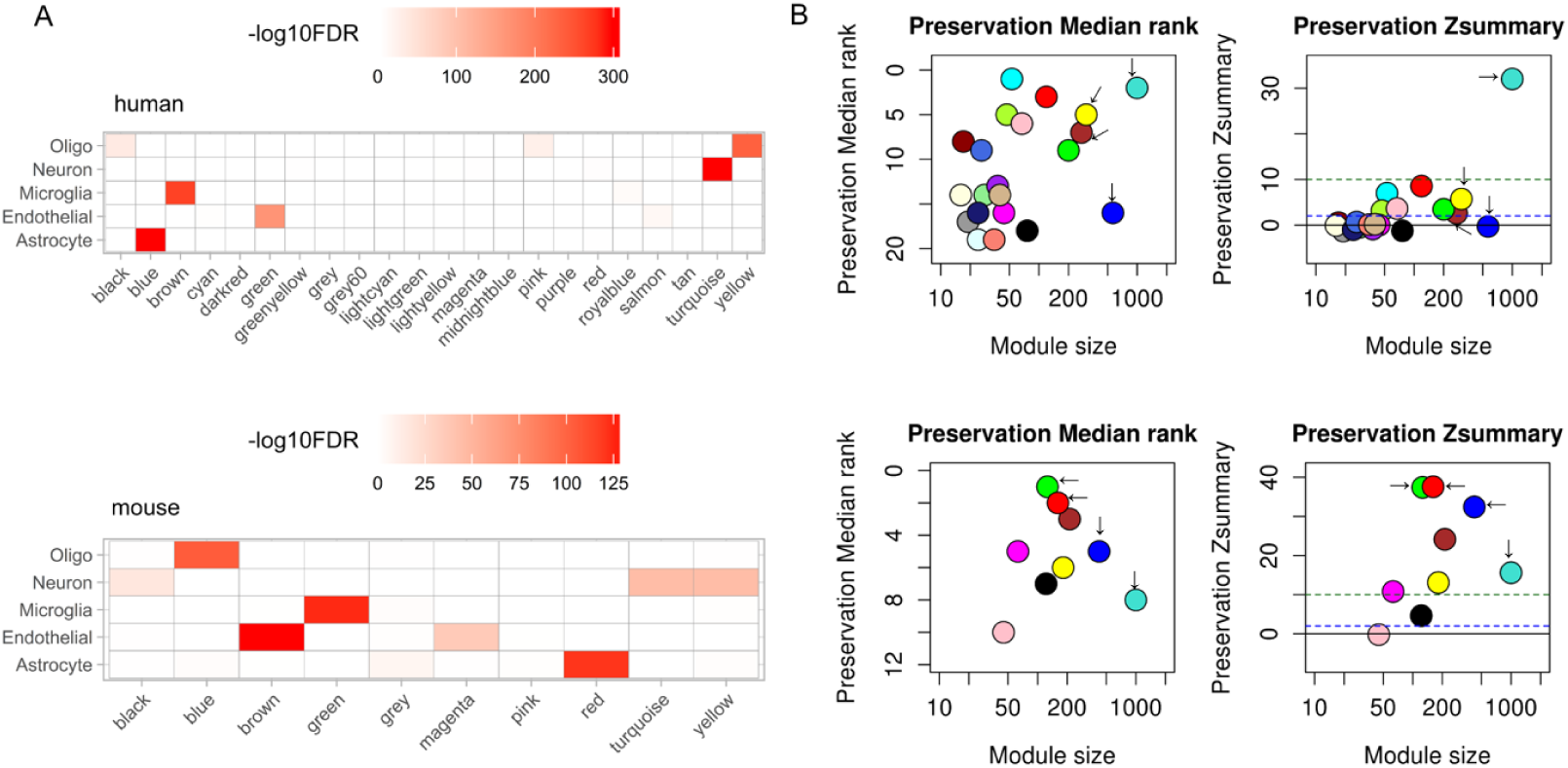
Cell type enrichment and preservation test of co-expression modules for human and mouse brain. (A) Enrichment of brain cell marker genes in human and mouse co-expression modules. The most significantly enriched module was defined as the brain cell co-expression module (BCCM) for each cell type. The human BCCMs are blue (astrocyte), brown (microglia), turquoise (neuron), and yellow (oligodendrocyte). The mouse BCCMs are red (astrocyte), green (microglia), turquoise (neuron), blue (oligodendrocyte). (B) Preservation of BCCMs between human and mouse brain. The top panel is the preservation test of BCCMs of the human brain in mouse data. The bottom panel is the preservation test of BCCMs of the mouse brain in human data. The arrows point to the BCCMs. Zsummary < 2 indicates the module is not preserved, 2 < Zsummary < 10 indicates weak to moderate preservation, and Zsummary > 10 indicates high module preservation.

**Table 3.**
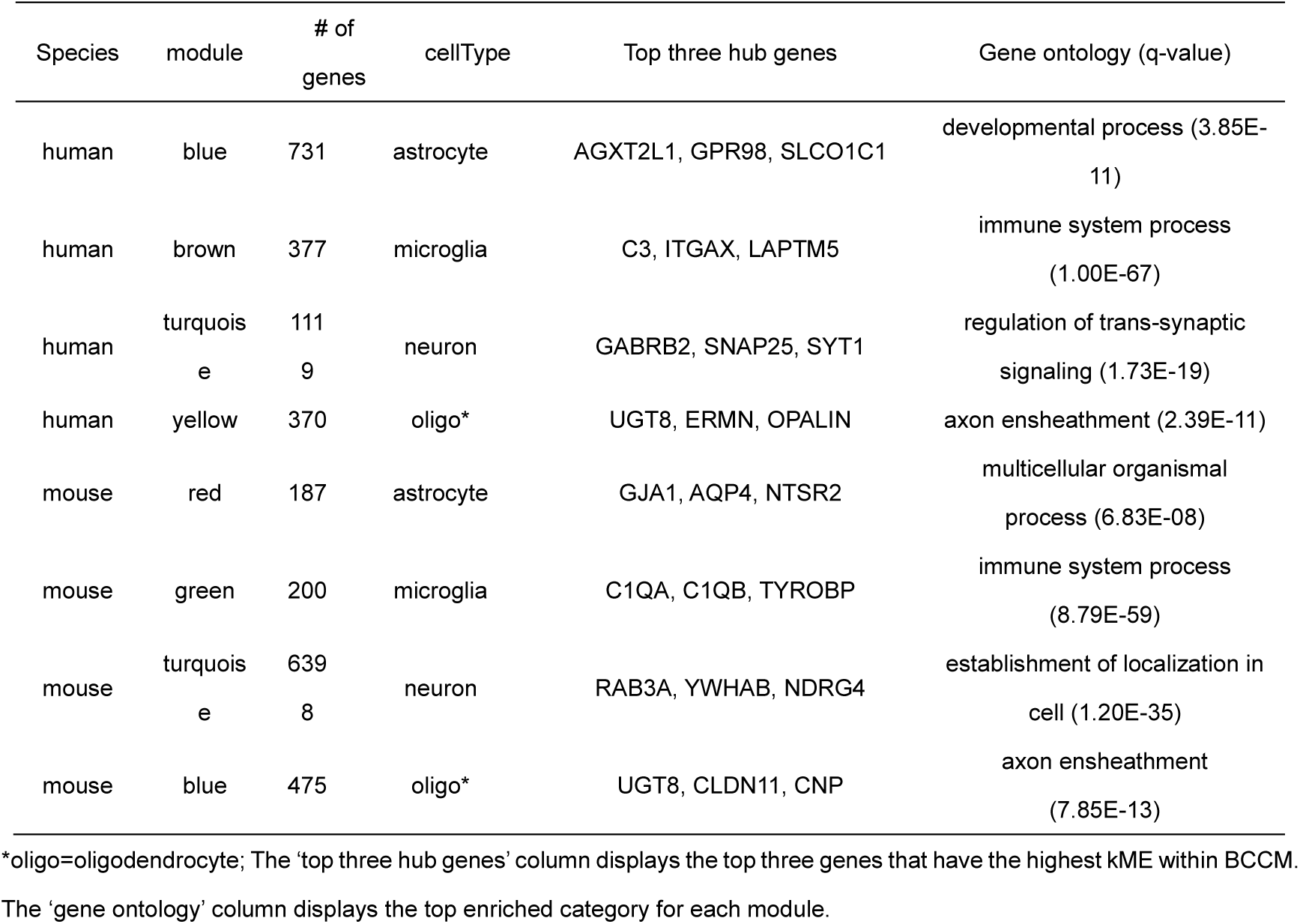
Brain cell co-expression modules in human and mouse

Next, we used the module preservation test to compare the BCCMs in human and mouse. The BCCMs of mouse brain were preserved in the human brain co-expression network. However, only the human neuron module was preserved in the mouse brain co-expression network (Figure 2B). Therefore, we analyzed the BCCMs for mouse and human brain separately in subsequent analysis to ensure we discover marker genes tailored specifically for human and mouse.

### DGE-COE relationship of brain cell marker genes

After the independent analyses of DGE and COE, we explored the relationships between them. We first asked whether marker genes with stronger specificity have a higher probability to enter the BCCMs than those with lower specificity. We tested 107 marker genes covered by six DGEDats and human COEDat. These 107 genes had 72 clustered into the four cell-type specific BCCMs and 35 into the other non-BCCMs. We found that csFC values of the 72 BCCM marker genes were higher than those of the 35 non-BCCM marker genes in all six DGEDats (Figure 3A, p-value of two-sample Wilcoxon test <0.05). In other words, marker genes in the BCCMs were more specific than the marker genes in the non-BCCMs. Significantly higher csFC values of marker genes in BCCMs than in non-BCCMs were also observed in mouse data (Supplementary Figure 5A, p-value of two-sample Wilcoxon test <0.05). This suggests that the highly-specific marker genes are more likely to be placed in a BCCM.

**Figure 3.**
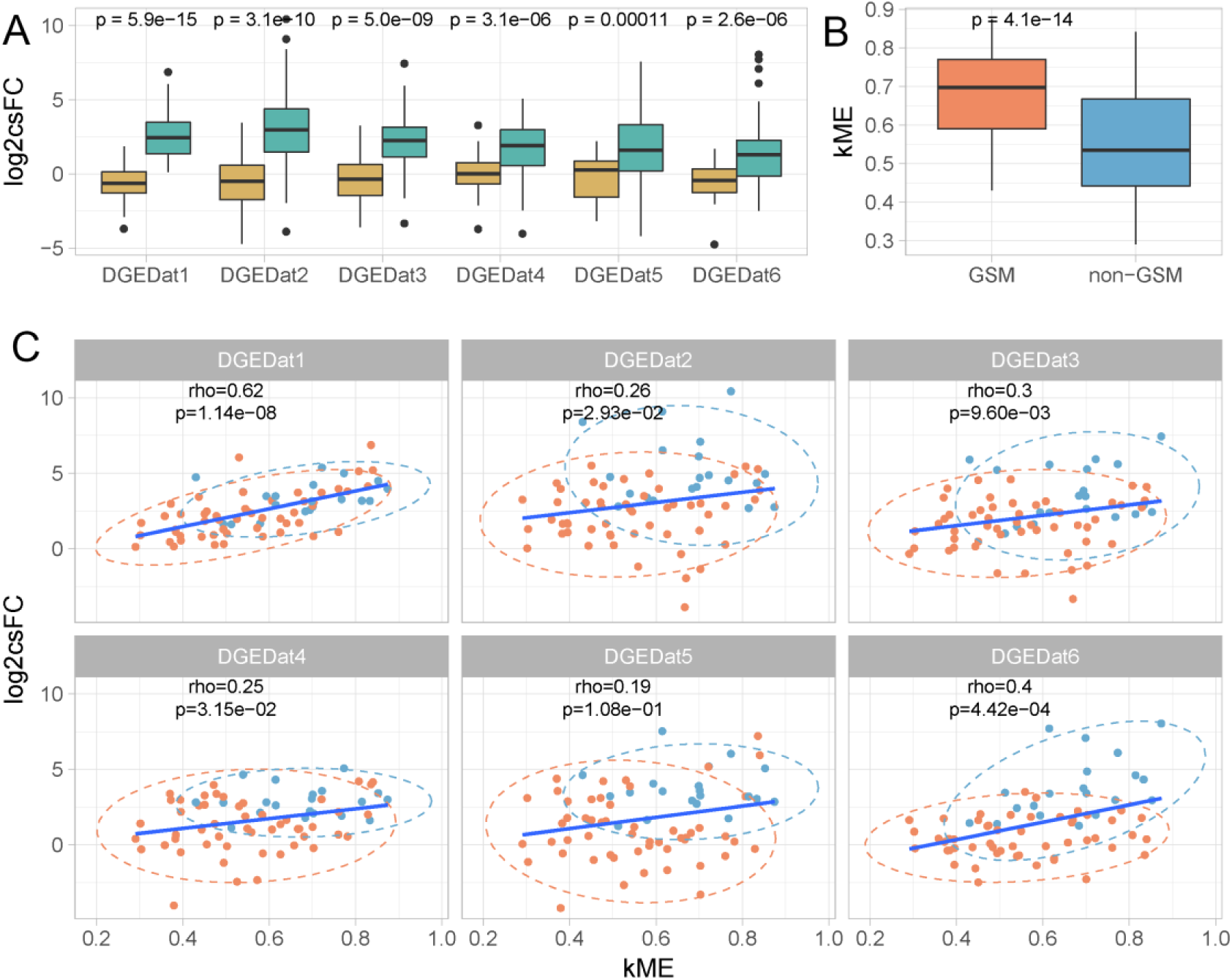
The relationship between DGE and COE of marker genes in human brains. (A) The comparison of csFC of BCCM marker genes and non-BCCM marker genes. The turquoise box denotes the marker genes in BCCMs and the mustard box denotes the marker genes in non-BCCMs (N_BCCM_ = 72, N_NON-BCCM_ = 35). The p-value is from a two-sample Wilcoxon test between csFC of marker genes in BCCMs and non-BCCMs. (B) The comparison of kME of the GSM and non-GSM in the BCCMs. A two-sample Wilcoxon test was used to test the significance of the difference (N_GSM_=20, N_non-GSM_=52). (C) The Spearman correlation between csFC and kME of marker genes in BCCMs. The blue dot represents GSM and the orange dot represent other marker genes. What are the dashed blue and orange circles?

Based on the test above, we next hypothesized that the highly-specific marker genes positioned close to the hub of the BCCMs have module membership rankings that are higher than non-GSM in the same BCCM. We divided the 72 marker genes in the human BCCMs into 20 GSM as identified above and 52 non-GSM. To compare the module membership ranking of these two gene groups, we performed a two-sample Wilcoxon test on their module membership (kME). kME is a measurement parameter used to assess the correlation between a gene and the eigengene, the hub of the co-expression module. A gene with high kME means that it has high correlations with other genes and consequently high ranking in the module. The kME values of GSM were significantly higher than those of non-GSM in the human BCCMs (p-value of two-sample Wilcoxon test<0.05, Figure 3B). However, the ranking of GSM in the BCCMs was not significantly higher than non-GSM in the mouse data (p-value of two-sample Wilcoxon test = 0.13, Supplementary Figure 5B).

These two analyses suggested that a connection did exist between DGE and COE for the marker genes. We further chose csFC representing DGE, and kME representing COE, to study the relationship between them. Significant correlations were observed between csFC values from five of the six DGEDats and kME values from human co-expression network (Spearman rho>0.2, p < 0.05; Figure 3C). In the mouse data, kME values of the marker genes were significantly correlated with csFC values in four of the six DGEDats (Spearman rho>0.2, p < 0.05; Supplementary Figure 5C). This indicates that high cell-specific fold change and high correlation with other marker genes in the BCCMs are two related properties of marker genes.

### Novel candidate brain cell marker genes are revealed by integration of COE and DGE

Based on the relationship observed between DGE and COE, we developed new criteria for selecting novel candidate brain cell marker genes (NCM). Since the BCCMs of human and mouse were not completely preserved, NCM was defined in human and mouse separately. The mouse NCM should have 1) csFC equal to or greater than 2 in at least two DGEDats from DGEDat2-DGEDat6 (BH corrected p-value of two samples of Wilcox test < 0.05), and 2) kME should be greater than 0.6 in COEDat2. We identified 16 mouse NCMs according to the criteria (Table 4, Supplementary Table S4). Because only one DGEDat for the human brain was available for analysis, we set relatively stricter criteria for human NCM to make more conservative calls. The human NCM should have 1) csFC significantly larger than 4 in the DGEDat1 (BH corrected p-value < 0.05) and 2) kME should be greater than 0.8 in the COEDat1. We identified 18 human NCM meeting these criteria (Table 4, Supplementary Table S5).

**Table 4.** NCM of human and mouse brain and their cellular locations

| gene | cellType | species | ISH | location |
| --- | --- | --- | --- | --- |
| ABCC9 | oligodendrocyte | human | - | Plasma membrane |
| ACSS1 | oligodendrocyte | human | - | Mitochondrial matrix |
| AHCYL1 | oligodendrocyte | human | - | Cytoplasm |
| CXCR7 | oligodendrocyte | human | - | Plasma membrane |
| DDAH1 | oligodendrocyte | human | - | Cytosol |
| EMX2OS | oligodendrocyte | human | - | - |
| GNA14 | oligodendrocyte | human | - | Plasma membrane |
| GPR125 | oligodendrocyte | human | - | Plasma membrane |
| IL33 | oligodendrocyte | human | - | Nucleoplasm |
| LRRC16A | oligodendrocyte | human | - | Plasma membrane |
| MT3 | oligodendrocyte | human | - | Nucleus |
| PAPLN | oligodendrocyte | human | - | Extracellular region |
| RHOJ | oligodendrocyte | human | - | Plasma membrane |
| SLC14A1 | oligodendrocyte | human | - | Plasma membrane |
| SNTA1 | astrocyte | human | Y | Plasma membrane |
| TIMP3 | astrocyte | human | - | Extracellular region |
| TPD52L1 | astrocyte | human | - | Cytoplasm |
| WIF1 | astrocyte | human | - | Extracellular region |
| C1qb | microglia | mouse | Y | Extracellular region |
| Mrc1 | microglia | mouse | Y | Plasma membrane |
| Csf1r | microglia | mouse | Y | Plasma membrane |
| Ctss | microglia | mouse | Y | Lysosome |
| Ptpn6 | microglia | mouse | Y | Nucleus |
| Cacna2d1 | neuron | mouse | Y | Plasma membrane |
| Elavl4 | neuron | mouse | - | Nucleus |
| SPin1 | neuron | mouse | Y | Nucleus |
| Gria1 | neuron | mouse | Y | Plasma membrane |
| Nipsnap1 | neuron | mouse | Y | Mitochondrion |
| Slc25a22 | neuron | mouse | Y | Plasma membrane |
| Mapk8 | neuron | mouse | Y | Nucleus |
| Stau2 | neuron | mouse | Y | Nucleus |
| Sirt2 | oligodendrocyte | mouse | Y | Nucleus |
| Bcas1 | oligodendrocyte | mouse | Y | Nucleus |
| Plxnb3 | oligodendrocyte | mouse | Y | Plasma membrane |

ISH: in situ hybridization image data from Allen Brain Atlas, Y: yes, having ISH image to confirm the locations, -: no ISH image.

### GSM and NCM improve the performance of supervised deconvolution

We used supervised deconvolution to examine how the choice of marker genes impacts deconvolution results using mouse data. We hypothesized that including GSM and NCM would improve deconvolution accuracy compared to not having them in the calculations. We downloaded mouse expression data from purified neuron, astrocyte, oligodendrocyte, and microglia, as well as RNA mixtures with known proportions of each cell type^31^. The purified cell expression data was used as a reference profile, and the mixture data was used for deconvolution. We constructed four types of reference gene sets: baseline, GSM_plus, NCM_plus, and NCM_GSM_plus. The baseline reference gene set included all the genes except for GSM and NCM. The other references were constructed by adding GSM, NCM, and their combination into the baseline reference. We used the root mean square error (RMSE) between estimated cell proportions and the true proportion to evaluate deconvolution performance. Higher RMSE indicated poorer performance of deconvolution. The optimal number of marker genes for deconvolution was determined (Materials and Methods). We found that the deconvolutions with baseline reference of 400 genes had the lowest RMSE, so we used this number of genes to construct the four tested references.

We observed that adding either set of GSM or NCM into the reference reduced the RMSE (Figure 4), suggesting that the inclusion of GSM and NCM can improve the performance of deconvolution. The reference including both NCM and GSM performed the best. To prove that the improved performance of the reference with NCM or GSM was not because of a larger number of marker genes used, we completed permutations by constructing three permutated references with randomly selected genes, excluding GSM and NCM. The permutation was repeated 1000 times for each type of permutated reference. Deconvolution using a reference with GSM or NCM outperformed the deconvolution using a permutated reference without GSM or NCM, showing that improved deconvolution performance when GSM and NCM were included was not related to the increased reference size (Figure 4B).

**Figure 4.**
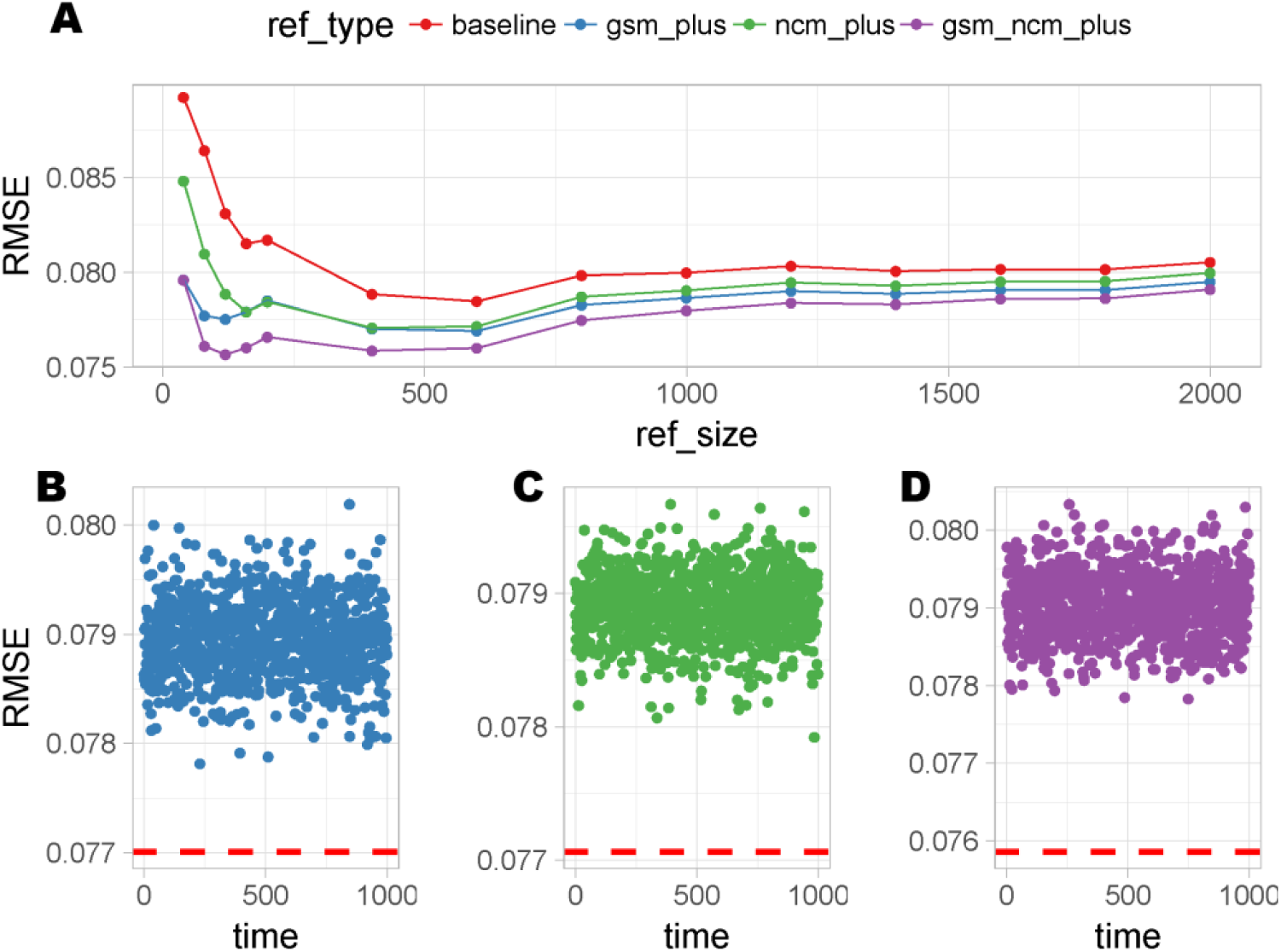
Effect of GSM and NCM in supervised deconvolution. (A) The RMSE between true and estimated cell proportion by supervised deconvolution with different references. The references are defined as follows: baseline = reference without GSM and mouse NCM; gsm_plus = baseline + GSM; ncm_plus = base + mouse NCM; gsm_ncm_plus = base + GSM +mouse NCM. With increasing size of the reference, the cell-specific fold change of marker genes included in the reference decreased. The deconvolution performance of permutated references without GSM and NCM where size is equal to the gsm_plus (B), ncm_plus (C), gsm_ncm_plus (D). The colors match the five refrences in figure 4A. The red dashed lines indicate the RMSE of deconvolution using gsm_plus, ncm_plus, and gsm_ncm_plus reference of 400 genes.

## Discussion

The current study describes the first systematic evaluation of marker gene specificity and their reliability for identifying cell types in human and mouse brains. We not only evaluated the published marker genes but also designed new criteria to discover novel marker genes based on both differential gene expression and co-expression. Applying our proposed novel marker genes to deconvolution improved the performance of deconvolution and resulted in more accurate cell proportion estimates.

This study identified a set of marker genes to discriminate neurons, astrocytes, microglia, and oligodendrocytes. New brain cell types have recently been identified with the development of single-cell RNA sequencing^32^. The evaluation of marker genes for these new cell types cannot be achieved currently because the multi-omics for these new cell types are not available. We required the cell types in evaluation to be measured at both transcriptome and proteome level, and currently only the four major cell types above satisfied the criteria. Our method will be adaptable to the newly identified brain cell types when multi-omics data are available.

One of the important outcomes of the current study was validating the specificity of marker genes reported in the literature. Most of the genes (304/540) included in the current study were claimed to be marker genes in a single source, and only eight genes had a consistent claim supported by all the collection sources (Supplementary Figure 1). Some genes that we tested (27 / 540) had conflict definitions for different cell types including several well-known marker genes, such as GFAP_^33^_ and ITGAM^34^. Our evaluation refined a list of reliable marker genes and supported using GFAP as a marker of astrocytes and ITGAM as a marker of microglia.

We were strict in assessing the specificity of marker genes, which led to removing some genes from commonly used marker gene lists. We compared the classic fold-change and cell type-specific fold-change of consistent marker genes (Supplementary Table 2). Eight marker genes were imprecisely defined in more than three of six DGEDats using the classic fold change. For example, SELENBP1 was a claimed astrocyte marker gene using averaged ranks across comparisons with each of other cell types^13^. However, its expression in microglia is close to, or even higher than expression in astrocytes in DGEDat2-DGEDat6. We removed it from the marker gene list because of its similar expression in microglia and astrocyte (Supplementary Figure 2). Most of the candidate marker genes failed to meet our criteria of GSM due to either being expressed at a similar level in more than two cell types (17%) or not being detectable as protein in the target cell type (20%), such as RBFOX3 and TMEM119. These two genes both showed target cell specificity when they could be detected (Supplementary Table 6). We expect that more marker genes including these two genes may be reclassified as GSM when more reliable proteomics data becomes available.

We showed a positive correlation between the csFC and kME of marker genes in both human and mouse brain. This is in line with our expectation that good marker genes will have similar expression patterns across cell types and strongly correlate with each other, which forms the core part of the cell module. The most important meaning of the strong correlation is that it suggests COE can be used for discovering marker genes. COE used all cell types, both characterized and uncharacterized, in brain tissue while DGE only used the several measured cell types to identify marker genes. The marker genes identified by COE should be more robust because they showed cell type-specificity across a broader range of cell types. This relationship will help to identify more brain cell marker genes from single-cell sequencing data, a technique that is increasing in popularity.

To explore the potential use of antibodies of NCM for cell labeling, we checked NCM’s subcellular localization of expression in the COMPARTMENTS database^35^ and the Allen Brain Atlas^36^. Eight human NCM and six mouse NCM are expressed on the plasma membrane, suggesting that antibodies made to these gene products have potential for use in FACS. One human NCM and seven mouse NCM are expressed at the nucleus, suggesting their potential use in sorting nuclei. Most of the mouse NCM already had archive ISH data except Elavl4. However, for the human brain, only SNTA1 had ISH data in the database. More experiments are needed to verify the subcellular location of the human NCM.

Supervised deconvolution was developed to replace the physical sorting of cell types. Supervised deconvolution infers cell proportion based on the expression of cell marker genes. Consequently, cell-type specificity of marker genes determines the accuracy of estimated proportions^37^. The deconvolution method is relatively well established, but validated marker genes for supervised deconvolution are lacking. NCM we proposed reduced the RMSE of deconvolution from 7.9% to 7.6% and resulted in improved accuracy of cell proportion estimates. The marginal improvement was expected because the baseline reference was composed of 400 genes with > 2-fold csFC. Instead of completing computations with 400 genes, using only the 21 GSM and 13 NCM we identified improved the performance of deconvolution slightly (0.3%) and is less resource intensive.

To date, various studies have found similarities and differences between tissue of humans and mice at the transcriptome level^17,38-40^. A study found a high degree of co-expression module preservation between human and mouse brain, and all mouse modules showed preservation with at least one human module whereas there were multiple human-specific modules^41^. The modules enriched in neuronal markers were more preserved between species than modules enriched glial marker genes^41^. This work conducted at the tissue level is consistent with our results showing that mouse shared BCCMs with human, but the BCCMs of the human brain were human-specific, except the neuron-related module. Our results also supported a recently published work at the single-cell level by Xu *et al.* who observed that hundreds of orthologous gene differences between human and rodent were cell type-specific^42^. Our data add to accumulating evidence that human have more cell-specific co-expression modules than mouse. Importantly, this implies that research on brain-related diseases using mouse models may have limited applicability to humans because of the difference between human and mouse brain cells. Furthermore, the definitions of brain cell types should consider species differences.

Our work is limited by the lack of cell-specific gene expression data with a large sample size and replication. This made the criteria for the evaluation less universal and more specific to our data sets. We could only calculate the p-value for four of six DGEDats due to lack of replication. Another limitation is the data used in the discovery of the relationship between DGE and COE were not from the same samples. This may explain why we did not observe strong correlations in all tested datasets.

Through a comprehensive evaluation of the brain cell marker genes; we developed a new method to identify marker genes, and provide a list of reliable marker genes for brain cells to guide the cell identification. Recently, studies reported methylome^43^ and regulome^44^ of brain cells, creating the potential to develop marker genes at epigenetics level. It would be meaningful to construct a framework by combining different omics data and methods to fully describe the cell types in the brain.

## Materials and Methods

### DGEdats pre-processing and quality control

We collected six datasets for the DGE-based evaluation. 1) DGEDat1^15^: Cells were isolated from the human temporal lobe cortex by immunopanning. We downloaded the fragments per kilobase of transcript per million mapped reads (FPKM) matrix. Fetal samples and genes with FPKM<0.1 in more than one sample were removed. 2) DGEDat2^14^: Cells were isolated from mouse cerebral cortex by immunopanning and FAC. We downloaded the expression level estimation which was quantified as FPKM. Genes with FPKM<0.1 in more than two samples were removed. 3) DGEDat3^10^: Gene expression of cells isolated from mouse brain cortex were measured by microarray. The microarray data contained 12 cell populations, which made use of the Mouse430v2 Affymetrix platform. We downloaded the raw CEL file. All the CEL files were subjected together to background correction, normalization and summary value calculation using the R package affy^45^ (‘rma’ function). The probes with ‘A’ or ‘M’ state in more than two samples were removed. 4) DGEDat4^11^: Cells were isolated from E16.5 and P1 mouse brain to culture neuron and glia cells. We downloaded the expression matrix which were quantified as reads per kilobase of transcript per million mapped reads (RPKM). Genes with RPKM<0.1 in more than three samples were removed. 5) DGEDat5 and DGEDat6^11^: both primary cultured cells and acutely isolated cells were collected from four replicates of 9-week-old whole mouse brains. Liquid chromatography-tandem mass spectrometry analysis was performed. We downloaded the quantified expression matrix. Genes with one missing value were removed.

### COEdats pre-processing and quality control

Two large-scale single-cell RNA sequencing datasets from both human and mouse brain were collected for co-expression analysis. 1) COEDat1. The human single cell transcriptome was from adult human individual’s temporal lobes^46^. In total, 332 cells from eight adult human brains (three males and five females) were collected and profiled by Illumina MiSeq and Illumina NextSeq 500. Raw sequencing reads were aligned using STAR and per gene counts were calculated using HTSEQ. We downloaded the counts matrix. 2) COEDat2. The mouse single cell transcriptomes of 3005 cells from somatosensory cortex and hippocampal CA1 regions were collected from juvenile (P22 - P32) CD1 mice including 33 males and 34 females^47^. The sequencing platform was Illumina HiSeq 2000. Raw reads were mapped to the mouse genome using Bowtie and the mapped reads were quantified to raw counts. We downloaded the counts matrix.

COEDats were pre-processed in Automated Single-cell Analysis Pipeline (ASAP)^48^. Genes with Counts per Million (CPM) lower than 1 in more than ten samples were removed from human brain data, and genes with CPM lower than 1 in more than 50 samples were removed from mouse brain data. After quality control, 13941 and 12149 genes were retained for human and mouse brain, respectively. The human brain data were normalized by voom function. Mouse data was normalized by scLVM. In total, 57 ERCC spike-ins in mouse data were used for fitting of technical noise. The normalized data were retained.

### Deconvolution data pre-processing and quality control

Gene expression data of brain samples with known cell proportion from rat was used in cell type-specific deconvolution^31^ (GEO accession: GSE19380). This dataset contains four different cell types including neuron, astrocyte oligodendrocyte and microglia, and two replicates of five different mixing proportions (Supplementary Table 7). The platform used was Affymetrix Rat Genome 230 2.0 Array. All the CEL files were subjected together to background correction, normalization and summary value calculation using ‘rma’ function.

### Co-expression analysis

To determine the gene networks of specific cell types, we completed weighted gene co-expression network analysis (WGCNA^22^) on single-cell sequencing data from both human and mouse brain using the signed network type. The parameter settings were as follows: Pearson correlation function, signed Topological Overlap Matrix (TOM) matrix, minimal module size of 20, deepSplit of 4, mergeCutHeight of 0.25 and pamStage of true. The power for human and mouse data was 7 and 6, respectively. The number of modules for human and mouse data was 22 and 10, respectively. The pSI package was used to identify the cell-related modules. The threshold for the enrichment test was BH-corrected p-value<0.05. The GO terms analysis was identified by Gorilla^49^. The expression localizations of genes were provided by COMPARTMENTS^35^.

### Module preservation test

A module preservation test was performed using the modulePreservation^50^ function in the WGCNA R package. Zsummary is a measurement to assess the preservation based on the size, density and the connectivity of modules. Zsummary < 2 indicated the module was not preserved, 2 < Zsummary < 10 indicated weak to moderate preservation, and Zsummary > 10 indicated high module preservation. We performed the module preservation test twice, once withmouse data as the reference and human data as the test set and once with roles reversed.

### Supervised deconvolution

We used function ‘lsfit’ in CellMix^4^ for deconvolution. In each mixture sample, we tested i probes and j cell types. The expression of each probe equals the sum of expression of purified cell types times corresponding cell proportions:

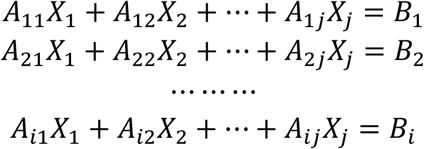

Where A_ij_ is an expression signal of probe i in a purified cell j, B_i_ is an expression signal of probe i in a mixture of cells, and X_j_ is a proportion of cell type j. The formula can be summarized in a matrix equation:

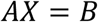

where A is the reference matrix of the expression of all probe sets in all cell types, B is the vector of expression levels of all probe sets in the mixture, and X is the vector of the proportions of all cell types comprising B. The equation was solved for X with the R function ‘lsfit’ (linear least squares algorithm).

The change of reference size was achieved by the following steps: 1) Construct the marker gene pool for four cell types and calculate the csFC. 2) Sort the marker gene pool according to the csFC in descending order. 3) Separate the reference genes into three types: GSM, NCM, and base genes. 4) Pick the desired number of marker genes from the base gene pool to construct baseline reference and perform deconvolution. 5) Add the GSM, mouse_NCM, or both GSM and NCM into the baseline reference to construct three tested references: gsm_plus, ncm_plus, gsm_ncm_plus. 6) perform deconvolution with three types of references separately. 7) Calculate RMSE between the estimated proportion and true proportion using the ‘rmse’ function in Metrics packages for each type of references. 9) Repeating step 2∼step 8 for increasing reference sizes.

## Acknowledgments

This work was supported by NSFC grants 81401114, 31571312, the National Key Plan for Scientific Research and Development of China (2016YFC1306000), and Innovation-Driven Project of Central South University (No. 2015CXS034, 2018CX033) (to C. Chen), and NIH grants 1U01 MH103340-01, 1R01ES024988 (to C. Liu). All the data contributors are sincerely thanked for the data provided. The authors thank Dr. Richard F. Kopp for critical reading of this manuscript.

## Author contributions

R.D. designed the study, performed the analyses and wrote the paper. Y.C., C.J., and J.D. helped with data collection and manuscript writing. C.L. and C.C created the project, supervised the study, contributed to the interpretation of the results, and revised the manuscript.

## Supplementary Materials

**Supplementary Table 1.** Collected commonly-used brain cell marker gene

**Supplementary Table 2.** The classical fold change and cell type-specific fold change of consistent marker gene

**Supplementary Table 3.** The GO term of BCCM for human and mouse

**Supplementary Table 4.** NCM of mouse brain cell

**Supplementary Table 5.** NCM of human brain cell

**Supplementary Table 6.** DGE of RBFOX3 and TMEM119

**Supplementary Table 7.** The true proportion of cell types in the mixture for deconvolution

**Supplementary Figure 1.**
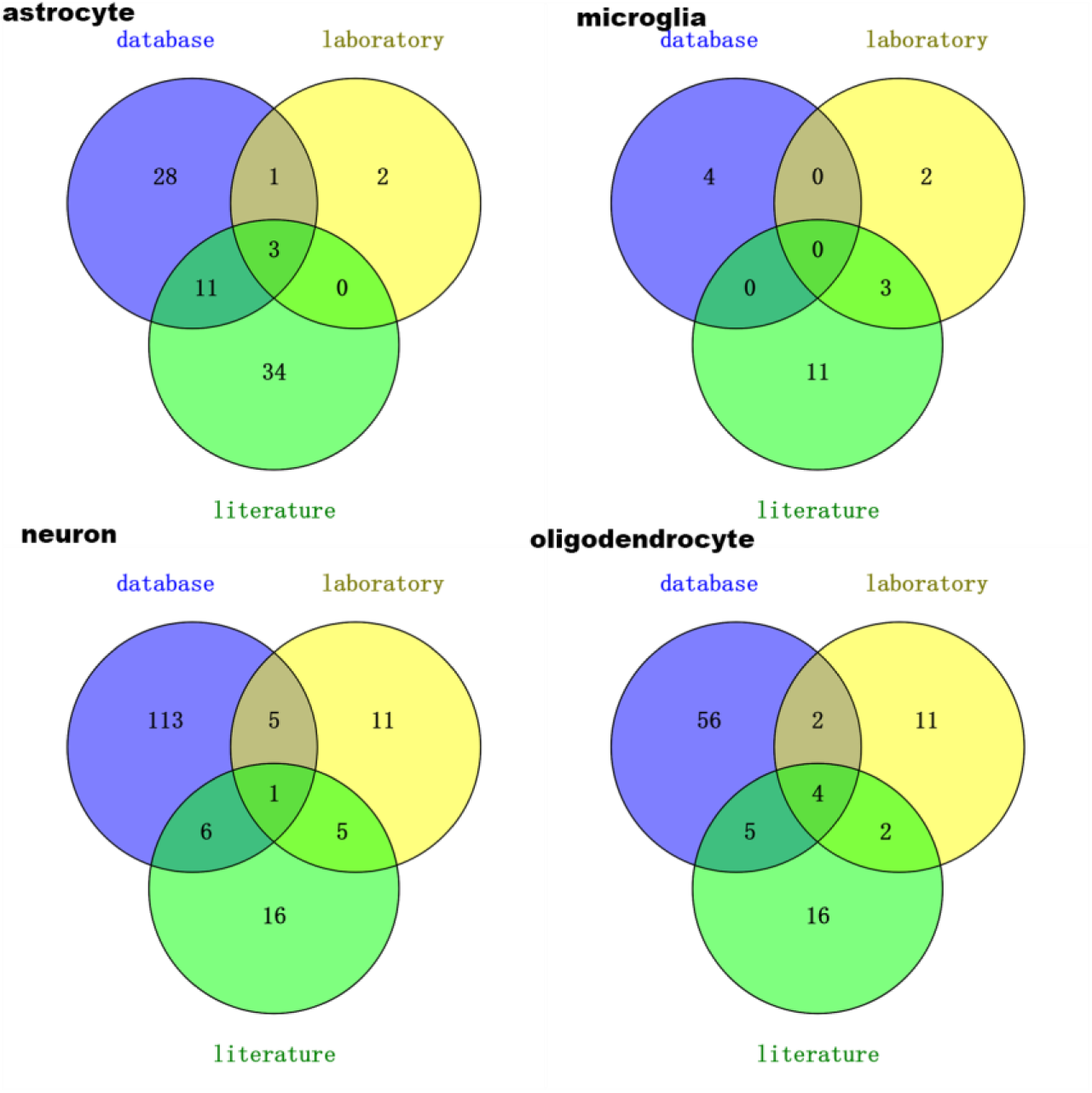
The overlap of marker genes collected from different sources. The commonly-used marker genes we evaluated were collected from three main sources: laboratory catalog, database, and published literature. The number indicates the number of marker genes belonging to corresponding sources.

**Supplementary Figure 2.**
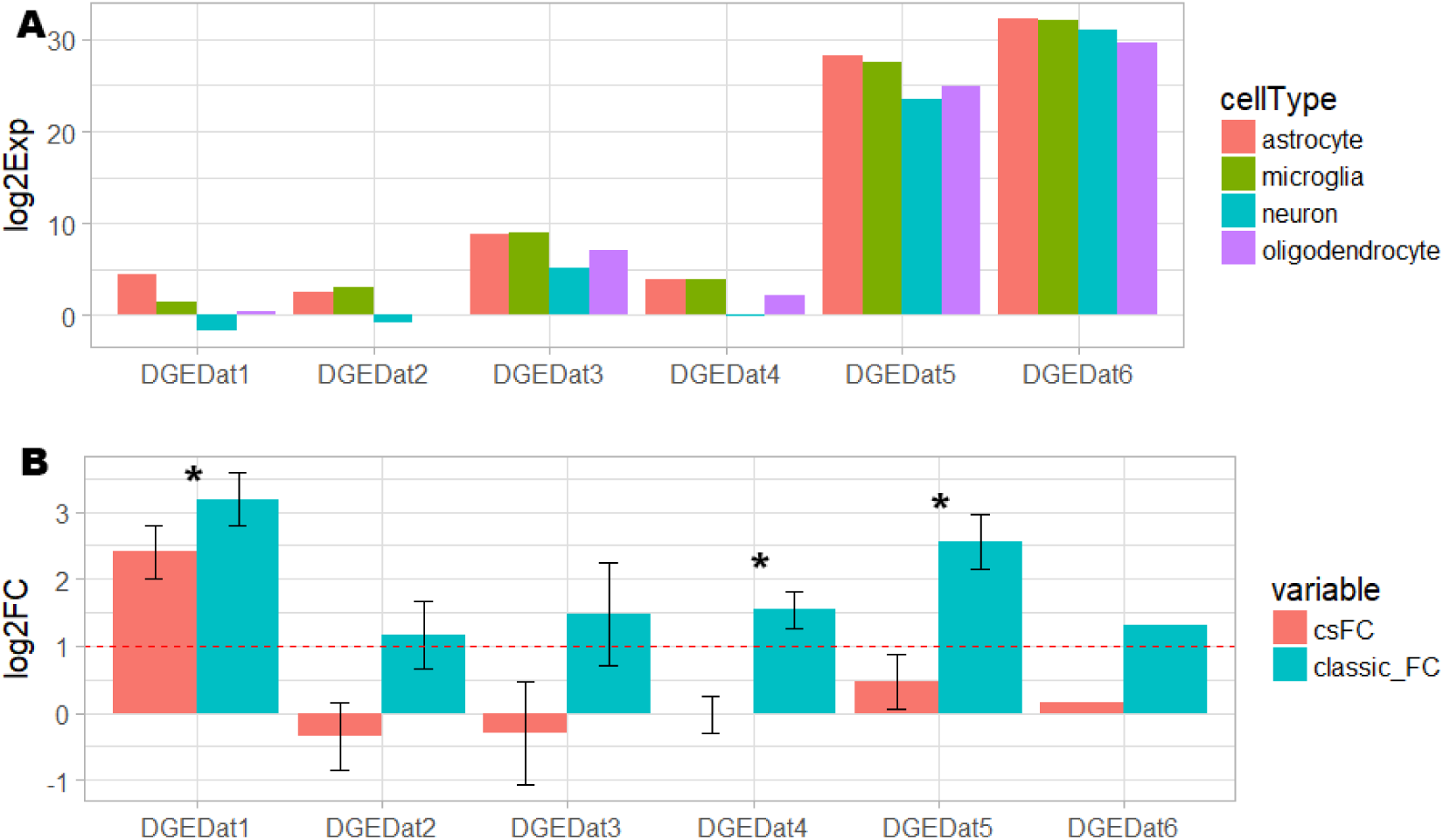
An example to illustrate the difference between cell-specific fold change and classical fold change. (A) The expression of SELENBP1. SELENBP1 is an un-validated marker gene of astrocyte. All six DGEDats detected it. Its expression in microglia is very close to even higher that the expression in astrocyte in DGEDat2-DGEDat6. (B) The fold change of SELENBP1. The cell type-specific fold change (csFC) and classical fold change for the SELENBP1 are measured. The red dashed line is the empirical cut-off for the fold change (log2FC=1). The error bar denotes the standard deviation of the fold change. The “*” indicate the BH-corrected p-value of two-sample Wilcoxon test is lower than 0.05. Since DGEDat6 have no replicates, the standard deviation cannot be calculated. The similar expression in the microglia will be covered up by the classical fold change calculation, while the csFC avoids this situation.

**Supplementary Figure 3.**
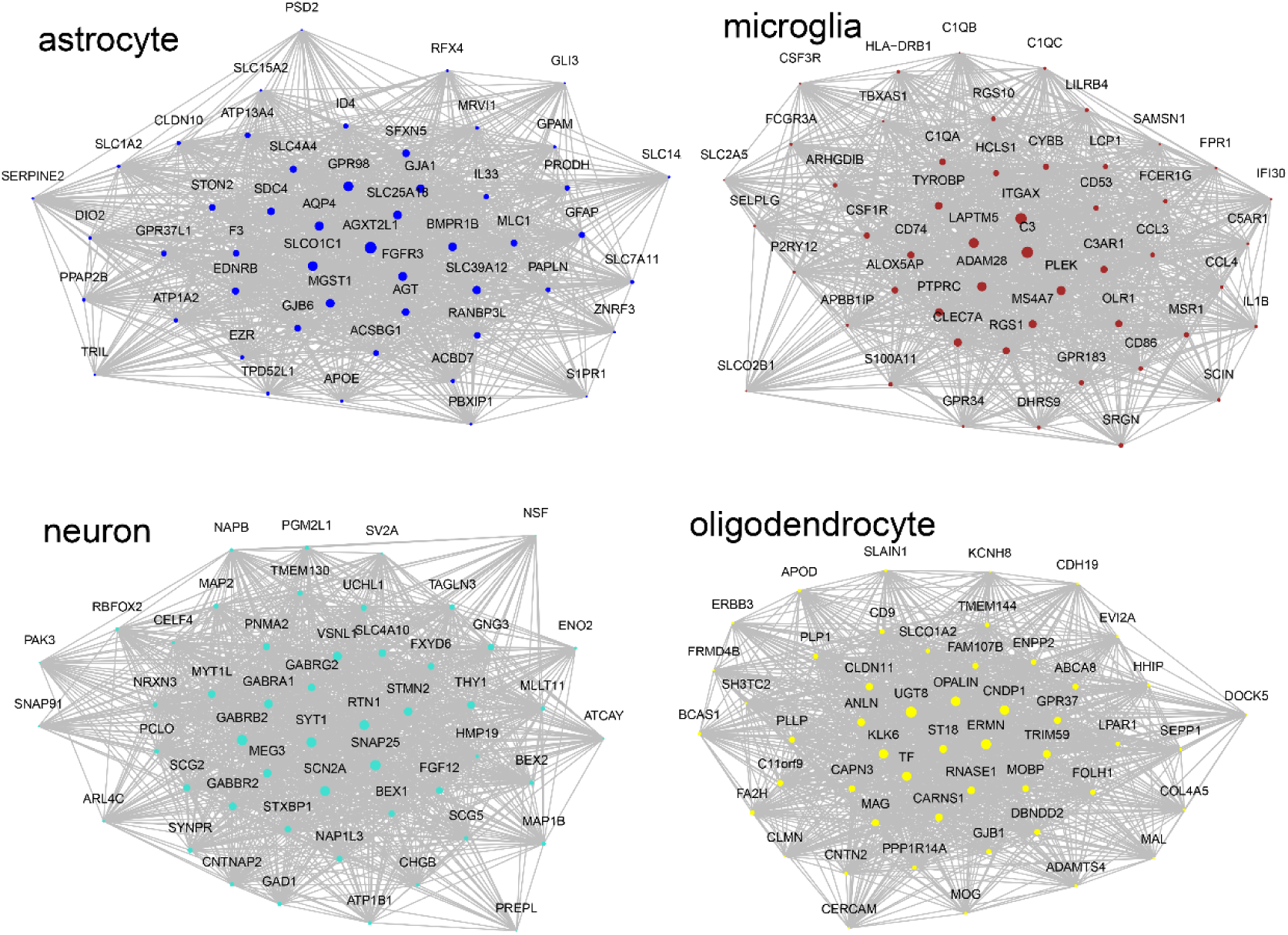
The top 50 hub genes of human brain cell co-expression module. The WGCNA was performed on human single-cell transcriptome. The brain cell co-expression module was selected according to the cell type enrichment conducted in pSI package. The gene members are ordered by kME from high to low. The dot color is the module color of brain cell co-expression module. The size of points indicates the kME of genes in the module with larger point representing higher kME.

**Supplementary Figure 4.**
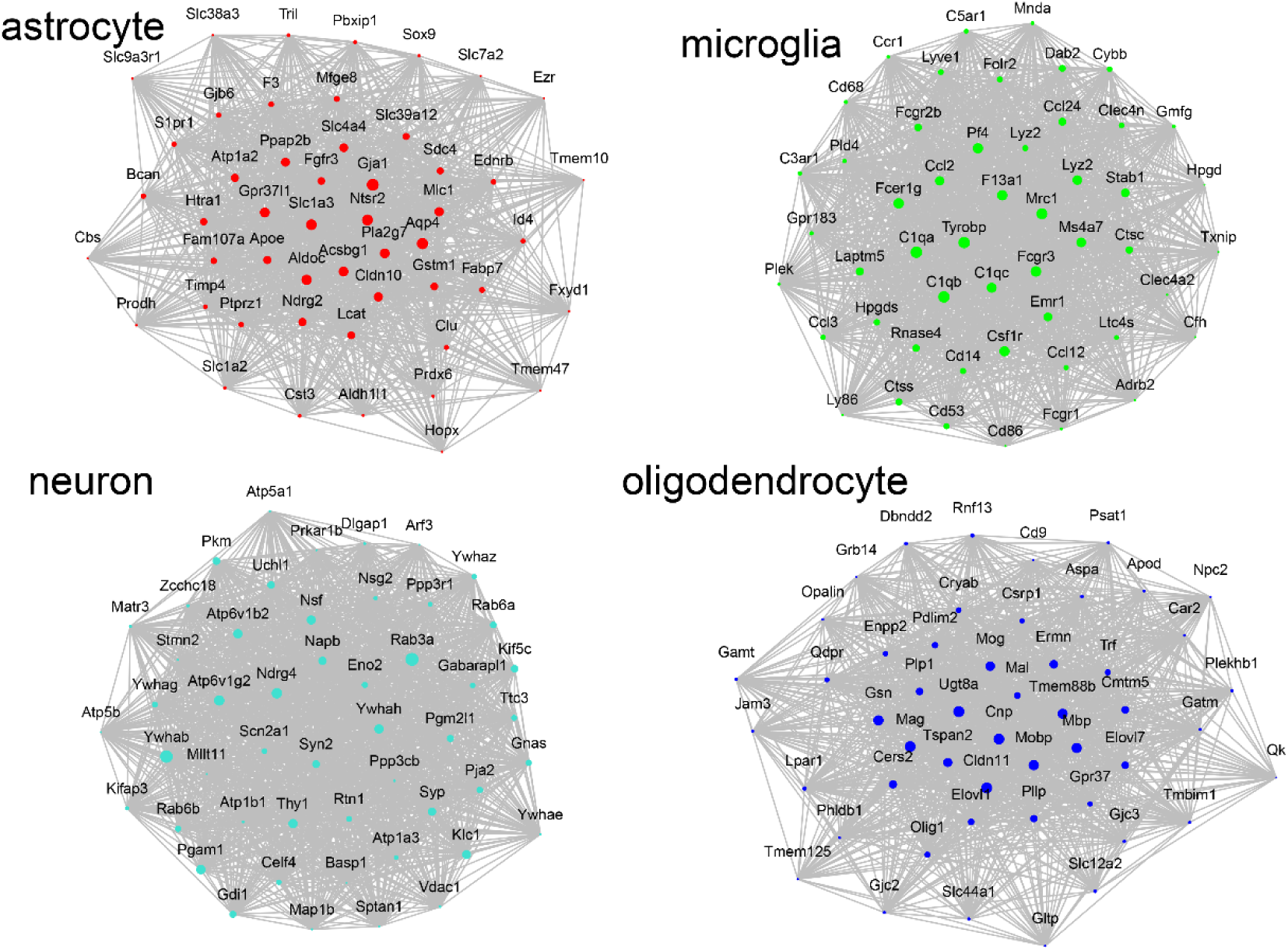
The top 50 hub genes of mouse brain cell co-expression module. The WGCNA was performed on mouse single-cell transcriptome. The brain cell co-expression module was selected according to the cell type enrichment conducted in pSI package. The gene members are ordered by kME from high to low. The dot color is the module color of brain cell co-expression module. The size of points indicates the kME of genes in the module with larger point representing higher kME.

**Supplementary Figure 5.**
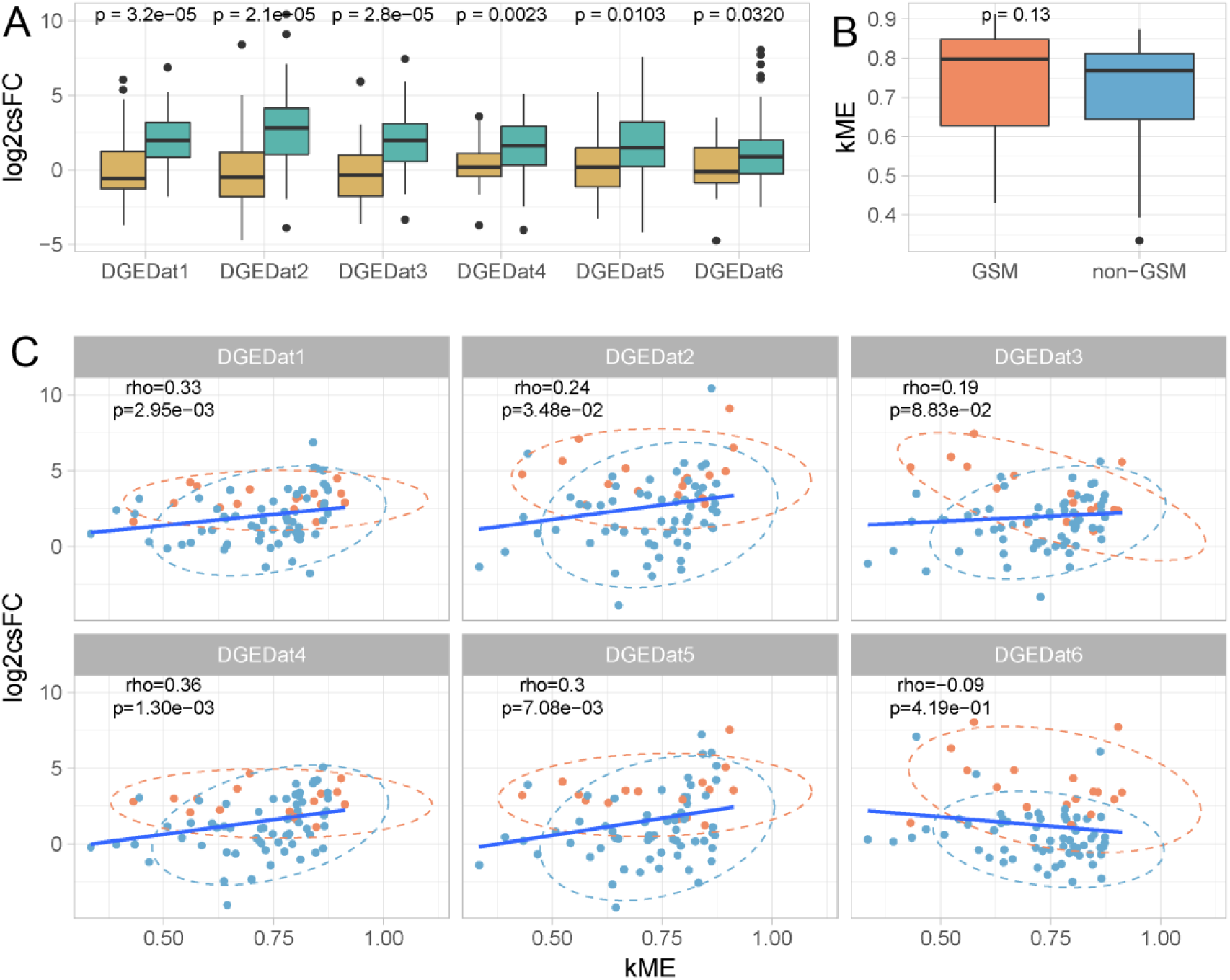
The relationship between DGE and COE in co-expression analysis of mouse data. (A) The comparison of csFC of brain cell co-expression module (BCCM) marker genes and non-BCCM marker genes. The turquoise box denotes the marker genes in BCCM and the mustard box denotes the marker genes in non-BCCM (N_BCCM_ = 79, N_NON-BCCM_ = 28). The p-value is from two-sample Wilcoxon test between csFC of marker genes in BCCMs and non-BCCMs. (B) The comparison of kME of the GSM and non-GSM in the BCCM. two-sample Wilcoxon test was used to test the significance of the difference (N_GSM_=19, N_non-GSM_=88). (C) The Spearman correlation between csFC and kME of marker genes in BCCMs. The blue dot represents GSM and the orange dot represent other marker genes.

**Supplemental Figure 6.**
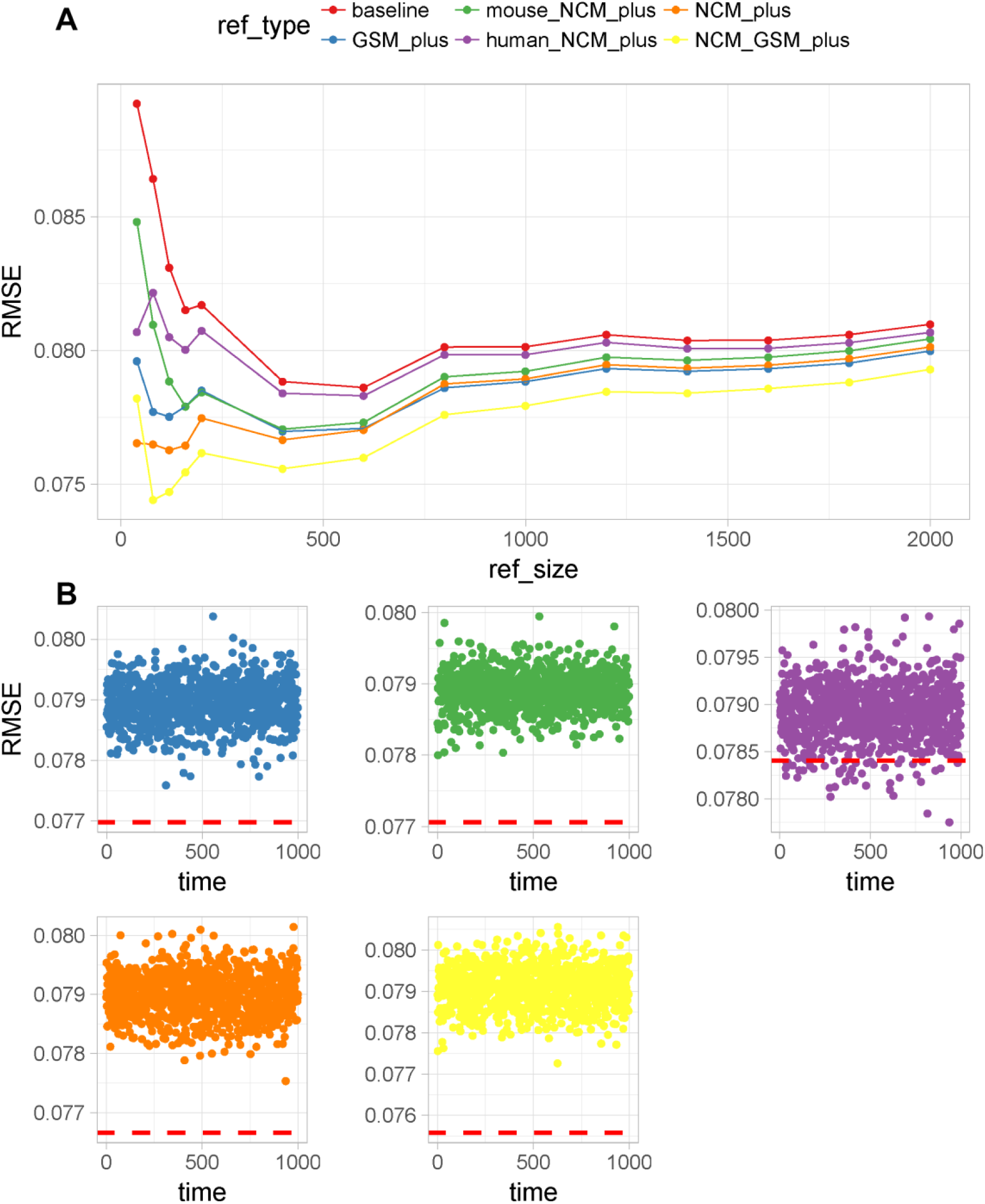
Effect of human GSM in deconvoluting mouse brain tissue. (A) The RMSE between true cell proportion and estimated cell proportion by supervised deconvolution with different references. The deconvolution performance of permutated references without GSM and NCM which size is equal to the reference tested above. The colors match the five references in figure 4A. The red dashed lines display the RMSE of deconvolution using tested reference of 400 genes.

## Reference

1 Azevedo, F. A. et al. Equal numbers of neuronal and nonneuronal cells make the human brain an isometrically scaled-up primate brain. The Journal of comparative neurology 513, 532–541, doi:10.1002/cne.21974 (2009).

2 Mancarci, B. O. et al. Cross-Laboratory Analysis of Brain Cell Type Transcriptomes with Applications to Interpretation of Bulk Tissue Data. eNeuro 4, doi:10.1523/ENEURO.0212-17.2017 (2017).

3 Mullen, R. J., Buck, C. R. & Smith, A. M. NeuN, a neuronal specific nuclear protein in vertebrates. Development 116, 201–211 (1992).

4 Gaujoux, R. & Seoighe, C. CellMix: a comprehensive toolbox for gene expression deconvolution. Bioinformatics 29, 2211–2212, doi:10.1093/bioinformatics/btt351 (2013).

5 Newman, A. M. et al. Robust enumeration of cell subsets from tissue expression profiles. Nature methods 12, 453–457, doi:10.1038/nmeth.3337 (2015).

6 Shen-Orr, S. S. & Gaujoux, R. Computational deconvolution: extracting cell type-specific information from heterogeneous samples. Current opinion in immunology 25, 571–578, doi:10.1016/j.coi.2013.09.015 (2013).

7 Fromer, M. et al. Gene expression elucidates functional impact of polygenic risk for schizophrenia. Nature neuroscience 19, 1442–1453, doi:10.1038/nn.4399 (2016).

8 Yu, Q. & He, Z. Comprehensive investigation of temporal and autism-associated cell type composition-dependent and independent gene expression changes in human brains. Scientific reports 7, 4121, doi:10.1038/s41598-017-04356-7 (2017).

9 Bachoo, R. M. et al. Molecular diversity of astrocytes with implications for neurological disorders. Proceedings of the National Academy of Sciences of the United States of America 101, 8384–8389, doi:10.1073/pnas.0402140101 (2004).

10 Cahoy, J. D. et al. A transcriptome database for astrocytes, neurons, and oligodendrocytes: a new resource for understanding brain development and function. The Journal of neuroscience : the official journal of the Society for Neuroscience 28, 264–278, doi:10.1523/JNEUROSCI.4178-07.2008 (2008).

11 Sharma, K. et al. Cell type- and brain region-resolved mouse brain proteome. Nature neuroscience 18, 1819–1831, doi:10.1038/nn.4160 (2015).

12 Sugino, K. et al. Molecular taxonomy of major neuronal classes in the adult mouse forebrain. Nature neuroscience 9, 99–107, doi:10.1038/nn1618 (2006).

13 Xu, X., Nehorai, A. & Dougherty, J. Cell Type Specific Analysis of Human Brain Transcriptome Data to Predict Alterations in Cellular Composition. Syst Biomed (Austin) 1, 151–160, doi:10.4161/sysb.25630 (2013).

14 Zhang, Y. et al. An RNA-sequencing transcriptome and splicing database of glia, neurons, and vascular cells of the cerebral cortex. The Journal of neuroscience : the official journal of the Society for Neuroscience 34, 11929–11947, doi:10.1523/JNEUROSCI.1860-14.2014 (2014).

15 Zhang, Y. et al. Purification and Characterization of Progenitor and Mature Human Astrocytes Reveals Transcriptional and Functional Differences with Mouse. Neuron 89, 37–53, doi:10.1016/j.neuron.2015.11.013 (2016).

16 Mouse Genome Sequencing, C. et al. Initial sequencing and comparative analysis of the mouse genome. Nature 420, 520–562, doi:10.1038/nature01262 (2002).

17 Lin, S. et al. Comparison of the transcriptional landscapes between human and mouse tissues. Proceedings of the National Academy of Sciences of the United States of America 111, 17224–17229, doi:10.1073/pnas.1413624111 (2014).

18 Januszyk, M. et al. Evaluating the Effect of Cell Culture on Gene Expression in Primary Tissue Samples Using Microfluidic-Based Single Cell Transcriptional Analysis. Microarrays 4, 540–550, doi:10.3390/microarrays4040540 (2015).

19 Schwanhausser, B. et al. Corrigendum: Global quantification of mammalian gene expression control. Nature 495, 126–127, doi:10.1038/nature11848 (2013).

20 Dong, X., You, Y. & Wu, J. Q. Building an RNA Sequencing Transcriptome of the Central Nervous System. Neuroscientist 22, 579–592, doi:10.1177/1073858415610541 (2016).

21 Zhang, B. & Horvath, S. A general framework for weighted gene co-expression network analysis. Statistical applications in genetics and molecular biology 4, Article17, doi:10.2202/1544-6115.1128 (2005).

22 Langfelder, P. & Horvath, S. WGCNA: an R package for weighted correlation network analysis. BMC bioinformatics 9, doi:10.1186/1471-2105-9-559 (2008).

23 Oldham, M. C. et al. Functional organization of the transcriptome in human brain. Nature neuroscience 11, 1271–1282, doi:10.1038/nn.2207 (2008).

24 abcam. <http://www.abcam.com/research/neuroscience/cell-type-marker> (

25 MerckMillipore. <http://www.merckmillipore.com/CN/zh/life-science-research/antibodies-assays/antibodies-overview/Research-Areas/neuroscience/Neurons-and-Glia/HtGb.qB.WxEAAAFPBc51gPtr,nav> (

26 Doyle, J. P. et al. Application of a translational profiling approach for the comparative analysis of CNS cell types. Cell 135, 749–762, doi:10.1016/j.cell.2008.10.029 (2008).

27 Lein, E. S. et al. Genome-wide atlas of gene expression in the adult mouse brain. Nature 445, 168–176, doi:10.1038/nature05453 (2007).

28 Hawrylycz, M. J. et al. An anatomically comprehensive atlas of the adult human brain transcriptome. Nature 489, 391–399, doi:10.1038/nature11405 (2012).

29 Dugas, J. C., Tai, Y. C., Speed, T. P., Ngai, J. & Barres, B. A. Functional genomic analysis of oligodendrocyte differentiation. The Journal of neuroscience : the official journal of the Society for Neuroscience 26, 10967–10983, doi:10.1523/JNEUROSCI.2572-06.2006 (2006).

30 Dougherty, J. D., Schmidt, E. F., Nakajima, M. & Heintz, N. Analytical approaches to RNA profiling data for the identification of genes enriched in specific cells. Nucleic acids research 38, 4218–4230, doi:10.1093/nar/gkq130 (2010).

31 Kuhn, A., Thu, D., Waldvogel, H. J., Faull, R. L. & Luthi-Carter, R. Population-specific expression analysis (PSEA) reveals molecular changes in diseased brain. Nature methods 8, 945–947, doi:10.1038/nmeth.1710 (2011).

32 Boldog, E. et al. Transcriptomic and morphophysiological evidence for a specialized human cortical GABAergic cell type. Nature neuroscience 21, 1185–1195, doi:10.1038/s41593-018-0205-2 (2018).

33 Eng, L. F., Ghirnikar, R. S. & Lee, Y. L. Glial fibrillary acidic protein: GFAP-thirty-one years (1969-2000). Neurochemical research 25, 1439–1451 (2000).

34 Cardona, A. E., Huang, D., Sasse, M. E. & Ransohoff, R. M. Isolation of murine microglial cells for RNA analysis or flow cytometry. Nature protocols 1, 1947–1951, doi:10.1038/nprot.2006.327 (2006).

35 Binder, J. X. et al. COMPARTMENTS: unification and visualization of protein subcellular localization evidence. Database : the journal of biological databases and curation 2014, bau012, doi:10.1093/database/bau012 (2014).

36 Sunkin, S. M. et al. Allen Brain Atlas: an integrated spatio-temporal portal for exploring the central nervous system. Nucleic acids research 41, D996-D1008, doi:10.1093/nar/gks1042 (2013).

37 Avila Cobos, F., Vandesompele, J., Mestdagh, P. & De Preter, K. Computational deconvolution of transcriptomics data from mixed cell populations. Bioinformatics 34, 1969–1979, doi:10.1093/bioinformatics/bty019 (2018).

38 Strand, A. D. et al. Conservation of regional gene expression in mouse and human brain. PLoS genetics 3, e59, doi:10.1371/journal.pgen.0030059 (2007).

39 Zheng-Bradley, X., Rung, J., Parkinson, H. & Brazma, A. Large scale comparison of global gene expression patterns in human and mouse. Genome biology 11, R124, doi:10.1186/gb-2010-11-12-r124 (2010).

40 Dowell, R. D. The similarity of gene expression between human and mouse tissues. Genome biology 12, 101, doi:10.1186/gb-2011-12-1-101 (2011).

41 Miller, J. A., Horvath, S. & Geschwind, D. H. Divergence of human and mouse brain transcriptome highlights Alzheimer disease pathways. Proceedings of the National Academy of Sciences of the United States of America 107, 12698–12703, doi:10.1073/pnas.0914257107 (2010).

42 Xu, X. et al. Species and cell-type properties of classically defined human and rodent neurons and glia. eLife 7, doi:10.7554/eLife.37551 (2018).

43 Luo, C. et al. Single-cell methylomes identify neuronal subtypes and regulatory elements in mammalian cortex. Science 357, 600–604, doi:10.1126/science.aan3351 (2017).

44 Lake, B. B. et al. Integrative single-cell analysis of transcriptional and epigenetic states in the human adult brain. Nature biotechnology 36, 70–80, doi:10.1038/nbt.4038 (2018).

45 Gautier, L., Cope, L., Bolstad, B. M. & Irizarry, R. A. affy--analysis of Affymetrix GeneChip data at the probe level. Bioinformatics 20, 307–315, doi:10.1093/bioinformatics/btg405 (2004).

46 Darmanis, S. et al. A survey of human brain transcriptome diversity at the single cell level. Proceedings of the National Academy of Sciences of the United States of America 112, 7285–7290, doi:10.1073/pnas.1507125112 (2015).

47 Zeisel, A. et al. Brain structure. Cell types in the mouse cortex and hippocampus revealed by single-cell RNA-seq. Science 347, 1138–1142, doi:10.1126/science.aaa1934 (2015).

48 Gardeux, V., David, F. P. A., Shajkofci, A., Schwalie, P. C. & Deplancke, B. ASAP: a web-based platform for the analysis and interactive visualization of single-cell RNA-seq data. Bioinformatics 33, 3123–3125, doi:10.1093/bioinformatics/btx337 (2017).

49 Eden, E., Navon, R., Steinfeld, I., Lipson, D. & Yakhini, Z. GOrilla: a tool for discovery and visualization of enriched GO terms in ranked gene lists. BMC bioinformatics 10, 48, doi:10.1186/1471-2105-10-48 (2009).

50 Langfelder, P., Luo, R., Oldham, M. C. & Horvath, S. Is my network module preserved and reproducible? PLoS computational biology 7, e1001057, doi:10.1371/journal.pcbi.1001057 (2011).

